# The functional roles of zebrafish *HoxA*- and *HoxD*-related clusters in the pectoral fin development

**DOI:** 10.1101/2024.07.05.600788

**Authors:** Mizuki Ishizaka, Akiteru Maeno, Hidemichi Nakazawa, Renka Fujii, Sae Oikawa, Taisei Tani, Haruna Kanno, Rina Koita, Akinori Kawamura

**Affiliations:** Division of Life Science, Graduate School of Science and Engineering, Saitama University, Shimo-Okubo 255, Sakura-ku, Saitama 338-8570, Japan; Cell Architecture Laboratory, National Institute of Genetics, Yata 1111, Mishima, Shizuoka 411-8540, Japan

**Keywords:** pectoral fin, *Hox* genes, zebrafish, X-ray CT scan

## Abstract

The paralogs 9-13 *Hox* genes in mouse *HoxA* and *HoxD* clusters are critical for limb development. When both *HoxA* and *HoxD* clusters are deleted in mice, significant limb truncation is observed compared to the phenotypes of single and compound mutants of *Hox9-13* genes in these clusters. In zebrafish, mutations in *hox13* genes in *HoxA*- and *HoxD*-related clusters result in abnormal morphology of pectoral fins, homologous to forelimbs. However, the effect of the simultaneous deletions of entire *HoxA*- and *HoxD*-related clusters on pectoral fin development remains unknown. Here, we generated mutants with several combinations of *hoxaa*, *hoxab*, and *hoxda* cluster deletions and analyzed the pectoral fin development. In *hoxaa^-/-^;hoxab^-/-^*;*hoxda^-/-^* larvae, the endoskeletal disc and the fin-fold are significantly shortened in developing pectoral fins. In addition, we show that this anomaly is due to defects in the pectoral fin growth after the fin bud formation. Furthermore, in the surviving adult mutants, micro-CT scanning reveals defects in the posterior portion of the pectoral fin which is thought to represent latent regions of the limb. Our results further support that the functional role of *HoxA* and *HoxD* clusters is conserved in the paired appendage formation in bony fishes.

## Introduction

In the development of bilaterian animals, *Hox* genes, which encode homeodomain transcription factors, play a crucial role in determining the positional identities along the body axis. *Hox* genes are typically organized in clusters known as *Hox* clusters. The order of the *Hox* genes within a cluster corresponds to their collinear expression patterns in developing embryos along the body axis^1–3^. During the early stages of vertebrate evolution, a single primitive *Hox* cluster, consisting of 1–13 paralogous groups, underwent two rounds of whole-genome duplication, resulting in quadruplication^4,5^. As a result, mice now possess four *Hox* clusters known as *HoxA*, *HoxB*, *HoxC*, and *HoxD*, each located on different chromosomes.

Loss-of-function studies using knockout mice have revealed that *Hox* genes play important roles in vertebrate development^6,7^. Specifically, the 9-13 paralogs of *Hox* genes in *HoxA* and *HoxD* clusters, which all correspond to homologous to the *Drosophila Abd-B* homeotic gene, have been shown to cooperatively contribute to limb development^8–15^. In mouse and chick, it has been observed that *Hoxa9* to *Hoxa13* and *Hoxd9* to *Hoxd13* genes exhibit nested and collinear expression domains in the mesenchymal cells of developing limbs^1,6,16^. This occurs through the successive activation of these *Hox* genes according to their positional order in *HoxA* and *HoxD* clusters. Furthermore, knockout mice for *Hoxa* and *Hoxd* genes have been generated, providing insights into the roles of these *Hoxa* and *Hoxd* genes in providing the regional identities of each segment of the limbs^8–15^. For instance, mice lacking *Hoxa9* and *Hoxd9* exhibit abnormalities in the stylopod^12^, which is the bone on the proximal side of the limb. Similarly, mice lacking *Hoxa13* and *Hoxd13* show defects in the autopod^13^, which is the bone on the distal side of the limb. Moreover, the simultaneous deletion of *HoxA* and *HoxD* clusters leads to severe truncation of forelimbs, particularly distal elements, in mice^14^.

Zebrafish possess seven *hox* clusters that were further duplicated through teleost-specific whole-genome duplication^17^. Currently, zebrafish have two clusters derived from *HoxA*, namely *hoxaa* and *hoxab.* On the other hand, only *hoxda* cluster, derived from *HoxD*, is present, as *hoxdb* cluster has been lost, except for a single microRNA^17^. In the formation of the pectoral fin, which is homologous to the forelimb, the 9-13 paralogous *hox* genes in zebrafish *hoxaa*, *hoxab*, and *hoxda* clusters exhibit similar collinear expression patterns to those of mouse and chick^18–20^. Loss-of-function study of *hoxa13a*, *hoxa13b*, and *hoxd13a* demonstrated the severe truncation of the pectoral fin in adult zebrafish^19^. These findings suggest a deep conserved functional homology of the posterior *HoxA-* and *HoxD*-related genes in the formation of the paired appendages in bony fishes. However, other posterior *hox* genes besides these *hox13* genes are also expressed during pectoral fin formation in zebrafish^18,20^, and the function of the entire *HoxA-* and *HoxD*-related clusters in pectoral fin development is yet to be elucidated.

In a previous study, we used the CRISPR-Cas9 system to generate mutants lacking each of the seven *hox* clusters in zebrafish^21^. Among the three *HoxA-* and *HoxD*-related clusters, only *hoxab* cluster deletion mutants showed shortening of the pectoral fin during embryogenesis^21^, suggesting functional redundancy among these *hox* clusters. In this study, we generated mutants with various combinations of deletions in *hoxaa*, *hoxab*, and *hoxda* clusters and examined the role of these clusters in pectoral fin development and skeletal structures of pectoral fins in adult fish.

## Results

Previous studies have shown that mouse *Hox* genes in *HoxA* and *HoxD* clusters cooperate to play essential roles in limb patterning^8–15^. The absence of all *HoxA* and *HoxD* functions results in a significant truncation of distal limb elements^14^. To investigate the role of zebrafish *hoxaa*, *hoxab*, and *hoxda* clusters, which are homologous to mouse *HoxA* and *HoxD* clusters, in zebrafish development, we focused on the pectoral fin, a homologous to tetrapod forelimb in this study. To analyze the pectoral fin phenotypes, we examined larvae derived from intercrosses between triple hemizygous mutants for *hoxaa*, *hoxab*, and *hoxda* clusters (Fig. 1A-H). At 3 dpf, we observed that triple homozygous deletion mutants (hereafter referred to *hoxaa^-/-^;hoxab^-/-^;hoxda^-/-^*) had present but severely shortened pectoral fins (Fig. 1H). The pectoral fin shortening in *hoxaa^-/-^;hoxab^-/-^;hoxda^-/-^* mutants was significantly greater than in any of the other *hox* cluster-deleted mutants. These results indicate that zebrafish *hoxaa*, *hoxab*, and *hoxda* clusters redundantly function in pectoral fin formation. The similar shortened pectoral fin/forelimbs phenotype in zebrafish and mouse *HoxA-* and *HoxD*-related mutants further suggest that the roles of *HoxA-* and *HoxD-*related clusters in paired appendage formation are conserved between zebrafish and mice.

**Figure 1.**
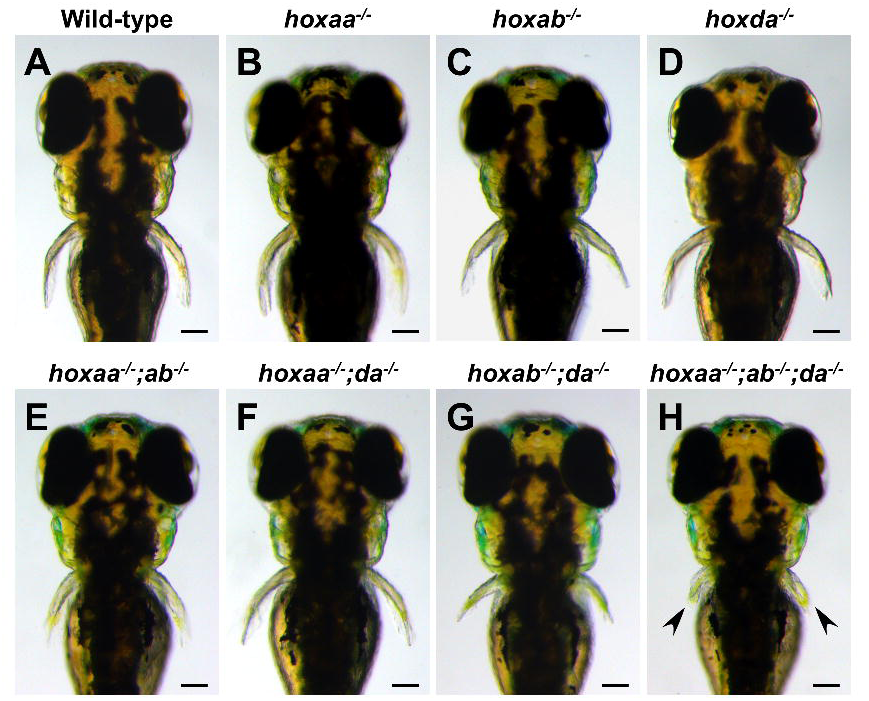
Shortening of pectoral fins in *hoxaa;hoxab*;*hoxda* cluster-deficient larvae. (A-H) Shortening of the pectoral fins in *hoxaa*-, *hoxab*, and *hoxda* cluster-deleted mutants. Dorsal views of larvae at 3 dpf. Arrowhead indicates severely truncated pectoral fins in *hoxaa*^-/-^;*hoxab*^-/-^ ;*hoxda*^-/-^ larva. Embryos were obtained from the natural intercrosses between *hoxaa*^+/-^ ;*hoxab*^+/-^ ;*hoxda*^+/-^ triple hemizygous mutant fish and were incubated at 28.5 C in water containing methylene blue as an anti-fungal agent (green signal in the otic vesicle). After photography, PCR-based genotyping was performed. At least three individuals (*n*>3) were identified for each genotype, and the representative larvae are shown. Scale bar, 100 μm.

The pectoral fin of zebrafish larvae consists mainly of the cartilaginous endoskeletal disc and the non-cartilaginous fin-fold (Fig. 2A)^22^. To examine the truncation of the pectoral fins in *hoxaa*, *hoxab*, and *hoxda* cluster mutant larvae in detail, we next compared the lengths of the endoskeletal disc and fin-fold, respectively (Fig. 2A-H). Cartilage-stained pectoral fins at 5 dpf showed that the lengths of the endoskeletal disc along the anterior-posterior and proximal-distal axes were significantly shorter in *hoxab^-/-^;hoxda^-/-^* and *hoxaa^-/-^;hoxab^-/-^;hoxda^-/-^* larvae, whereas the other combinations of *hoxaa*, *hoxab*, and *hoxda* cluster deletions did not appear to affect the lengths of the endoskeletal disc (Fig. 2I, J). On the other hand, the effects of *hoxaa*, *hoxab*, and *hoxda* cluster deletions were more pronounced with respect to the length of fin-fold (Fig. 2K). The fin-fold in *hoxaa^-/-^;hoxab^-/-^* larvae was shortened compared to that of wild-type larvae, whereas the differences in the length of the endoskeletal disc were not observed. Among the three double *hox* cluster deletion mutants, *hoxab^-/-^;hoxda^-/-^* larvae were the shortest in both the fin-fold and the endoskeletal disc. Furthermore, the fin-fold length was shortest in *hoxaa^-/-^;hoxab^-/-^;hoxda^-/-^* larvae, as was the endoskeletal disc. These results suggest that the cooperative functions of *hoxaa*, *hoxab*, and *hoxda* clusters: *hoxab* cluster has the highest contribution to pectoral fin formation in zebrafish, followed by *hoxda* cluster and then *hoxaa* cluster.

**Figure 2.**
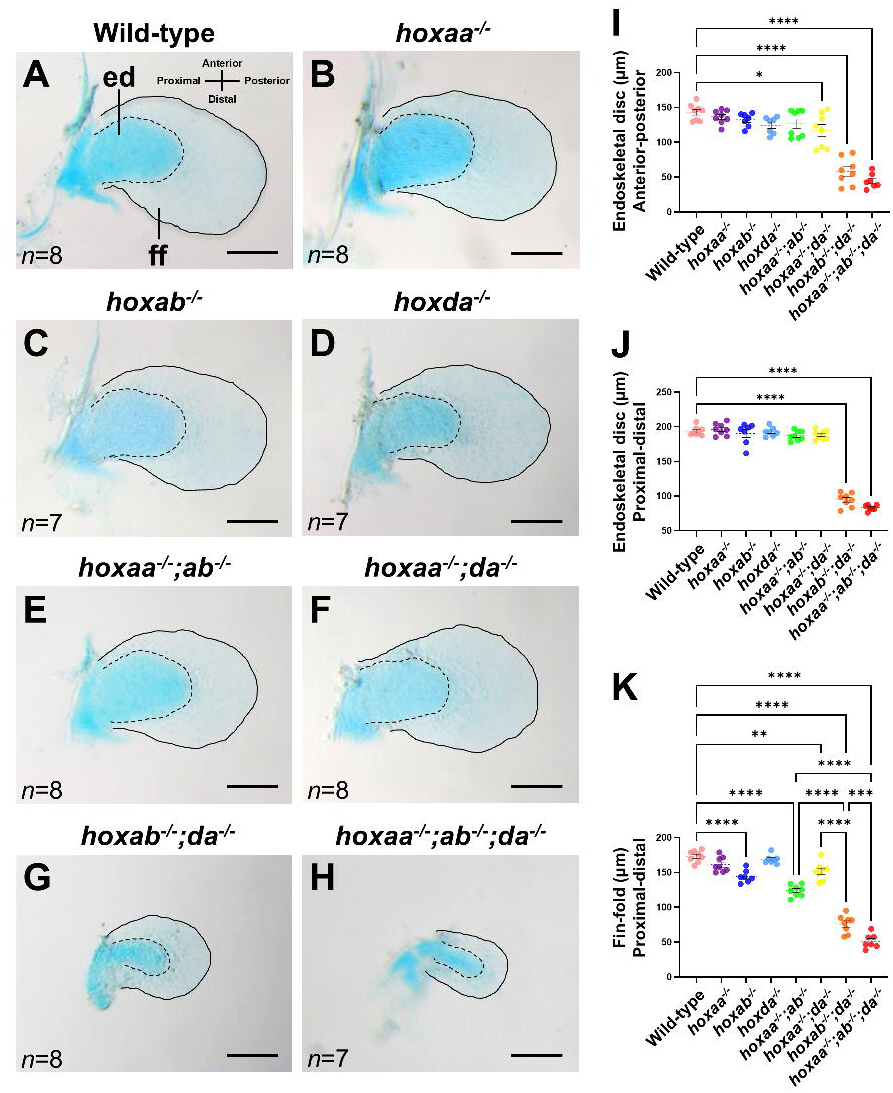
Shortening of the endoskeletal disc and fin-fold of the pectoral fins in hoxaa^-/-^ ;hoxab^-/-^ ;hoxda^-/-^ larvae. (A-H) Reduction of the endoskeletal disc (ed) and fin-fold (ff) in the pectoral fins of *hoxaa;hoxab;hoxda* cluster-deleted mutants. Alcian blue staining was performed to distinguish cartilaginous cells (endoskeletal disc) and non-cartilaginous fin-fold in larvae at 5 dpf. After the staining, the pectoral fins were manually dissected and mounted on a glass slide. The margin of the endoskeletal disc is emphasized with a dashed line, and the margin of the fin-fold is highlighted with a line. Proximal to the left. Scale bar, 100 μm. The number of dissected pectoral fins for which lengths were measured is noted in the left bottom. (I-K) Comparisons of the length of the endoskeletal disc and fin-fold in *hoxaa;hoxab;hoxda* cluster-deleted mutants. The lengths of the endoskeletal disc and the fin-fold were measured by ImageJ. The Tukey-Kramer test was performed with \**P*L<L0.05, \*\**P*L<L0.01, \*\*\**P*L<L0.001, and \*\*\*\**P*L<L0.0001. Error bars represent the standard error of the mean.

To gain further insight into the shortening of pectoral fins in *hoxaa;hoxab;hoxda* cluster mutants, we performed whole-mount *in situ* hybridization to examine the expression patterns of genes essential for the pectoral fin formation. Zebrafish *tbx5a* is essential for the initial formation of the pectoral fin buds, and loss of *tbx5a* function leads to the absence of the pectoral fins^23,24^. Thus, we examined the expression patterns of *tbx5a* in 30 hpf embryos derived from the intercrosses between triple hemizygous mutants for *hoxaa*, *hoxab*, and *hoxda* clusters. We found no significant differences in the expression of *tbx5a* in embryos obtained from this intercross. Furthermore, we confirmed by genotyping of the stained embryos that expression patterns of *tbx5a* in *hoxaa^-/-^;hoxab^-/-^*;*hoxda^-/-^*larvae appeared indistinguishable from that of sibling wild-type embryos (Fig. 3A, B). These results indicate the normal establishment of the pectoral fin buds even in the absence of all the functions of *hoxaa*, *hoxab*, and *hoxda* clusters. Therefore, we analyzed the expression of *sonic hedgehog a* (*shha*) (Fig. 3C-J), which is subsequently required for the cell proliferation of the developing pectoral fin^25^. In zebrafish, *shha* is expressed in the posterior portion of the fin buds in 48 hpf embryos (Fig. 3C). Consistent with shortened pectoral fins in zebrafish *HoxA-* and *HoxD*-related cluster mutants (Fig. 1, 2), reduced expression of *shha* in the pectoral fin buds was observed. This was particularly evident in *hoxab^-/-^*;*hoxda^-/-^* and *hoxaa^-/-^;hoxab^-/-^*;*hoxda^-/-^* larvae, where *shha* expression in fin buds was markedly down-regulated (Fig. 3I, J, Fig. S2). These results suggest that the shortening of the pectoral fin observed in zebrafish *HoxA-* and *HoxD*-related cluster mutants is due to retardation of pectoral fin growth after the normal formation of the fin buds by *tbx5a* expression.

**Figure 3.**
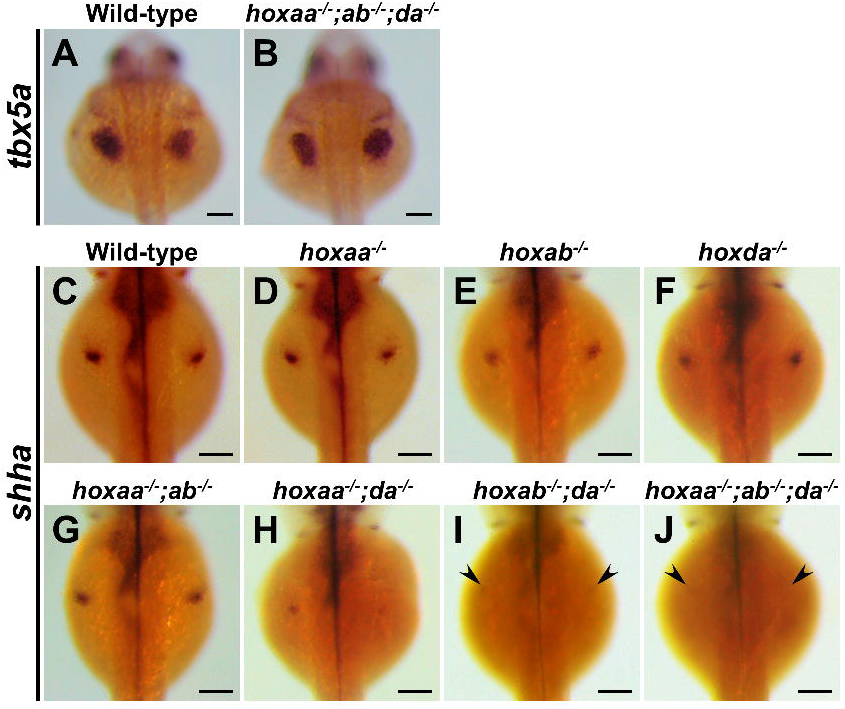
Expression of *shha* in developing pectoral fin is significantly decreased in *hoxaa;hoxab;hoxda* triple homozygous mutants. Whole-mount *in situ* hybridization was performed. Dorsal view. (A, B) Expression patterns of *tbx5a* in wild-type siblings (*n*=5) and *hoxaa*^-/-^ *;hoxab*^-/-^ *;hoxda*^-/-^ embryos (*n*=3) at 24 hpf. (C-J) Expression patterns of *shha* in the embryos at 48 hpf which were derived from several intercrosses between *hoxaa*^+/-^ *;hoxab*^+/-^ *;hoxda*^+/-^ fish. Prior to imaging, PCR-based genotyping was performed using genomic DNA extracted from the tail of the stained embryos. At least three embryos were identified for each genotype to confirm the reproducibility. Arrowhead indicates significantly decreased signal of *shha* in the pectoral fin primordium of *hoxab*^+/-^ *;hoxda*^+/-^ and *hoxaa*^+/-^ *;hoxab*^+/-^ *;hoxda*^+/-^ embryos. Scale bar, 100 μm.

Next, we investigated whether *HoxA-* and *HoxD-*related clusters contribute to the skeletal formation of adult zebrafish pectoral fins. For this purpose, we raised larvae to adult fish by multiple intercrosses between triple hemizygous mutants for *hoxaa*, *hoxab*, and *hoxda* clusters. Our previous study demonstrated that homozygous deletion of zebrafish *hoxab* cluster leads to lethality^21^. Consistently, fish with homozygous deletion in *hoxab* cluster did not survive with any combination of deletions in *hoxaa* and *hoxda* clusters (Table 1). Furthermore, fish with homozygous deletion of two or three clusters in *hoxaa*, *hoxab*, and *hoxda* also did not survive. Therefore, we analyzed the skeletal structures of the pectoral fins in *hoxaa^-/-^;hoxab^+/-^*;*hoxda^+/-^* and *hoxaa^+/-^;hoxab^+/-^*;*hoxda^-/-^* fish, which were the surviving adult fish with the most of these deletion mutations by using X-ray micro-CT scanning.

**Table 1. Genotyping of juvenile fish derived from the intercrosses between *hoxaa;hoxab;hoxda* triple hemizygous fish.**

Embryos were obtained by the natural spawning of intercrosses between triple hemizygous *hoxaa;hoxab;hoxda* fish and reared to juveniles of 1-2 months of age. Their genotype was then determined by PCR using genomic DNA extracted from the dissected caudal fin. The expected percentage was calculated according to Mendel’s law. It appears that the nonviable mutants die after 7 days after fertilization.

In our wild-type adult zebrafish, the number of fin rays in the pectoral fins is 10.8 on average (*n*=6, three specimens were examined for left and right pectoral fins; Fig. 4A, J, Table 2). By contrast, statistical analysis showed that the number of fin rays was slightly decreased to an average of 9.7 in the pectoral fins of *hoxaa^-/-^;hoxab^+/-^*;*hoxda^+/-^* fish (*n*=6; Fig. 4B, J, Table 2), whereas that of *hoxaa^+/-^;hoxab^+/-^*;*hoxda^-/-^* fish was comparable to that of wild-type fish (*n*=6; Fig. 4C, J, Table 2). In addition, we observed that the anterior fin rays formed normally as in the wild type, whereas one or two of the posterior fin rays were missing in *hoxaa^-/-^;hoxab^+/-^*;*hoxda^+/-^* fish (Fig. 4B, Table 2; *n*=6/6, 100 % affected). Furthermore, *hoxaa^-/-^;hoxab^+/-^*;*hoxda^+/-^* fish lost the pointed morphology of the proximal tips of the posterior 7th and/or 8th fin rays (Table 2; *n*=3/6, 50 % affected). *hoxaa^+/-^;hoxab^+/-^*;*hoxda ^-/-^* fish retained the normal number of fin rays, but also lost the pointed morphology of the proximal tips of the posterior four or five fin rays (Table 2; *n*=3/6, 50 % affected). On the other hand, in the radial domains of endoskeletons supporting the fin rays, the proximal radials are divided into four elements (pr1-pr4) along the anteroposterior axis in wild-type fish ^22^ (Fig. 4A,D), although we noticed that our wild-type zebrafish exhibited minor anomalies in the pectoral fins at low frequency by micro-CT scanning (Table 2). However, *hoxaa^-/-^;hoxab^+/-^*;*hoxda^+/-^* fish exhibited the absence of the most posteriorly located pr4 and instead showed an enlargement of pr3 (*n*=5/6, 83 % affected; Fig. 4B, Table 2). Similarly, *hoxaa^+/-^;hoxab^+/-^*;*hoxda^-/-^* fish also showed the absence of pr4 (*n*=5/6, 83 % affected; Fig. 4C,F,I, Table 2). Furthermore, we also observed a pectoral fin that lacked pr3 along with the absence of pr4 (*n*=3/6, 50 % affected; Fig. 4C, Table 2). Previous study revealed that zebrafish *hoxa13a^-/-^;a13b^-/-^* and *hoxa13a^-/-^;a13b^-/-^;d13a^-/-^* mosaic triple mutants showd the increased number of distal radials^19^. To clarify whether the number of distal radials is also increased in *hoxaa^-/-^;hoxab^+/-^*;*hoxda^+/-^*and *hoxaa^+/-^;hoxab^+/-^*;*hoxda^-/-^* fish, segmentation of micro-CT scan images was performed to visualize the distal radials. In the wild-type, the number of distal radials is 7.3 on average (Fig. 4G, K, Table 2). However, in *hoxaa^-/-^;hoxab^+/-^*;*hoxda^+/-^* fish, the number of the distal radials averaged 11.2, and in *hoxaa^+/-^;hoxab^+/-^*;*hoxda^-/-^* fish, the number increased to an average of 11.5 (Fig. 4H, I, K, Table 2). Comparing the degree of abnormalities in the pectoral fins, slightly stronger abnormalities were observed in *hoxaa^+/-^;hoxab^+/-^*;*hoxda^-/-^* fish than in *hoxaa^-/-^;hoxab^+/-^*;*hoxda^+/-^* fish. This result may reflect the high contribution of the *hoxda* cluster among the three *HoxA-* and *HoxD*-related clusters. These results suggest that in zebrafish mutants with reduced function of *hoxaa*, *hoxab*, and *hoxda* clusters, a pronounced structural defect was commonly observed in the posterior part of the pectoral fins such as pr3, pr4, and the tip of the posterior fin rays. Interestingly, the posterior part of the pectoral fin is thought to share molecular similarities with the limbs of tetrapods^18,26–31^. Thus, the present loss-of-function study further supports the notion that *Hox* genes in the *HoxA* and *HoxD* clusters already participated in patterning the paired appendages before the divergence of the ray-finned and lobe-finned fishes.

**Figure 4.**
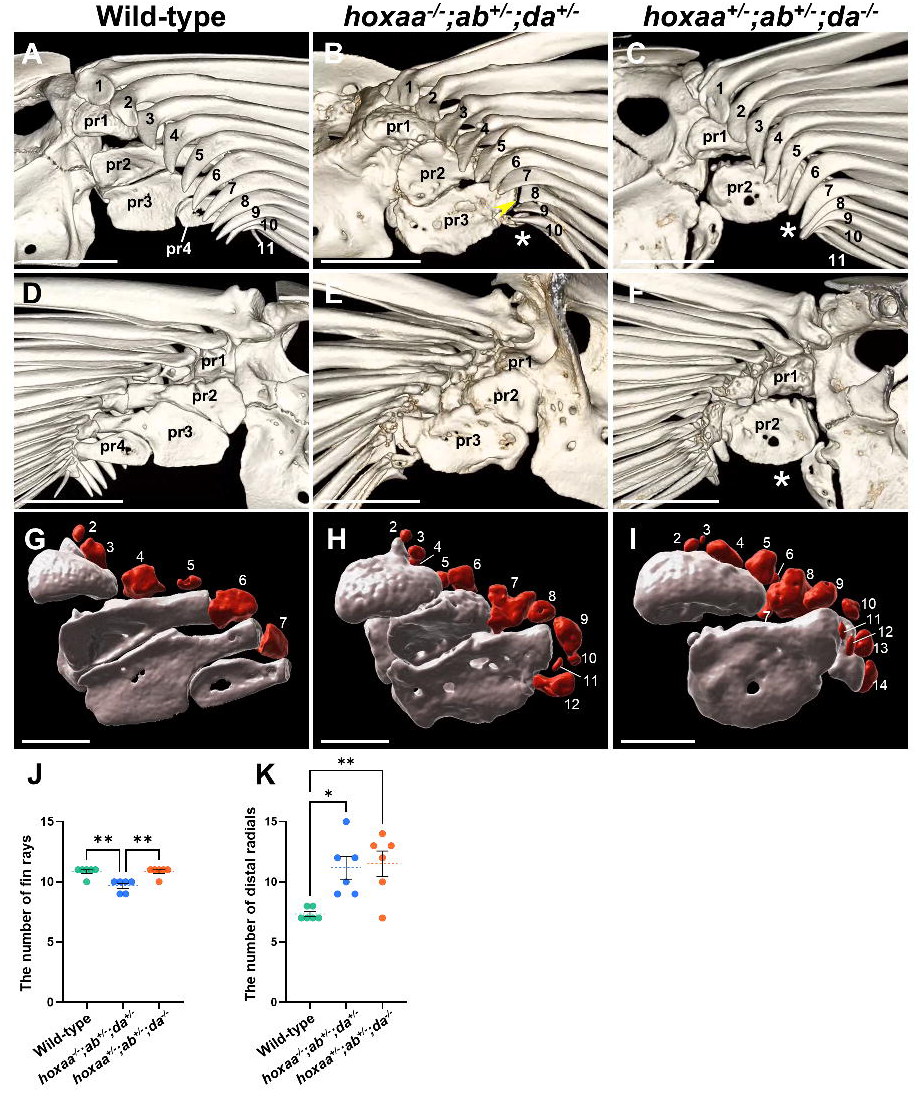
Skeletal elements in the posterior regions of the pectoral fin are preferentially affected in *hoxaa*^-/-^ *;hoxab*^+/-^ *;hoxda*^+/-^ and *hoxaa*^+/-^ *;hoxab*^+/-^ *;hoxda*^-/-^ adult fish. Micro-CT scan analysis of the skeletal structures in zebrafish pectoral fins. Both sides of the pectoral fins in wild-type (*n*=3), *hoxaa*^-/-^ *;hoxab*^+/-^ *;hoxda*^-+/-^ (*n*=3), and *hoxaa*^+/-^ *;hoxab*^+/-^ *;hoxda*^+/-^ (*n*=3) adult zebrafish were analyzed. The left pectoral fins are shown. (A-C) The magnified views of pectoral fin endoskeletons from the lateral side. (D-F) Magnified views of the pectoral fin endoskeletons from the opposite medial side in the same specimens shown in A-C. Proximal radials (pr1-pr4) are indicated. Fin rays are numbered from the anterior to the posterior. The asterisk indicates the absence of the posterior proximal radials. The arrowhead indicates the loss of the proximal tip of the fin rays. Anterior is to the top. The 3D movies were created using OsiriX MD ver13 and are available in Movies S1-3. Scale bar, 500 μm. (G-I) Magnified images of proximal radials (grey) and distal radials (red) as revealed by CT segmentation. The same specimens as above were used for each genotype. Fin rays were removed by image processing. The first distal radials are not shown due to their distance from the remaining distal radials. 3D movies were created using Imaris ver10 (Movies S4-6). A 2D movie showing the distal radials and proximal radials was also created using Imaris ver10 and is provided in Movie S7. Scale bar, 250 μm. (J,K) Comparisons of the number of fin rays and distal radials between wild-type, *hoxaa*^-/-^*;hoxab*^+/-^*;hoxda*^+/-^ and *hoxaa*^+/-^*;hoxab*^+/-^*;hoxda*^-/-^ adult fish. The Tukey-Kramer test was performed with \**P*L<L0.05 and \*\**P*L<L0.01.

**Table 2. Summary of the abnormal morphology of the skeletal elements in pectoral fins.**

The left and right pectoral fins of three specimens each of wild-type, *hoxaa^-/-^;hoxab^+/-^;hoxda^+/-^*, and *hoxaa^+/-^;hoxab^+/-^;hoxda^-/-^* mutants were analyzed by micro-CT scanning and the abnormalities in the skeletal elements were summarized. In the table, “-” indicates the normal morphology. For each genotype, #1 is shown in Fig. 4.

## Discussion

In this study, we generated compound mutants with different combinations of zebrafish *hoxaa*, *hoxab*, and *hoxda* cluster deletions and analyzed their role in the pectoral fin development. Combining these three *hox* cluster deletions increases early pectoral fin development severity, suggesting that *hox* genes in *hoxaa*, *hoxab*, and *hoxda* clusters function cooperatively as was observed in mouse *HoxA* and *HoxD* clusters^8–15^. In a previous study, zebrafish *hoxa13a;hoxa13b* double mutants were generated, showing that the endoskeletal disc is almost normal but the finfold is shortened by about 30 % in the pectoral fin of larvae^19^. In this study, we showed the pectoral fin phenotype of mutants lacking all *hoxaa*, *hoxab*, and *hoxda* clusters, and both endoskeletal disc and finfold were found to be significantly shortened in *hoxaa^-/-^;hoxab^-/-^*;*hoxda^-/-^* larvae. These abnormalities in our mutants are apparently stronger than the pectoral fin abnormalities observed in *hoxa13a;hoxa13b* double mutants^19^. The paralogous *hox* 9-13 genes of *HoxA-* and *HoxD*-related clusters are expressed overlappingly in the developing pectoral fin in zebrafish^18,20^, strongly suggesting that these *hox9*-*hox13* genes contribute to pectoral fin formation in a coordinated manner as was observed in mice.

When comparing the phenotypes of double *hox* cluster deletions in early pectoral fin formation, the combination of *hoxab* and *hoxda* cluster deletions causes the most severe phenotype (Fig. 1, 2). Detailed expression pattern analysis of the developing pectoral fin demonstrated that zebrafish *hox9-13* genes in *hoxab* and *hoxda* clusters are overlappingly expressed in mesenchymal cells in the fin buds from early stages. In contrast, *hox* genes in *hoxaa* cluster are not expressed early, but are expressed late in distal mesenchymal cells in the pectoral fins^18^. These similarities of the expression patterns support that the most severe phenotype of *hoxab^-/-^*;*hoxda^-/-^* larvae in pectoral fin formation. It is also possible that the combination of deletions in *hoxab* and *hoxda* clusters is the only combination of the three types of *hox* double deletions that cannot be functionally complemented in part because the remaining *hoxaa* cluster has no paralogous *hox10* and *hox12* (Fig. S1).

In the absence of 19 *hox* genes of zebrafish *hoxaa*, *hoxab*, and *hoxda* clusters, we observed that the expression of *tbx5a* in the pectoral fin buds was not altered (Fig. 3A, B). These results suggest that the formation of the pectoral fin buds occurs normally but subsequent cell differentiation or cell proliferation in the pectoral fin growth is severely impaired in *hoxaa^-/-^;hoxab^-/-^*;*hoxda^-/-^* larvae. Consistent with this observation, after the formation of fin buds, we found that *shha* expression in the developing pectoral fins is significantly down-regulated in mutants lacking zebrafish *HoxA-* and *HoxD*-related clusters (Fig. 3C-J). Since zebrafish *shha/syu* mutants have been shown to exhibit substantially reduced cell proliferation in pectoral fin buds which results in the truncation of pectoral fins^25,32^, the shortening of the pectoral fins in *hoxaa^-/-^;hoxab^-/-^*;*hoxda^-/-^* larvae could be primarily due to reduced *shha* expression in fin buds. In addition, these results are similar to previous results in mice that *Shh* expression was not detectable in the severely truncated forelimbs of *HoxA* and *HoxD* cluster double mutant mice^14^. Therefore, our loss-of-function results suggest not only that the functions of *Hox* genes in *HoxA* and *HoxD* clusters are conserved in pectoral fin and forelimb development, but also that the regulatory network comprising *Hox-Shh* genes in appendage formation was already established in common ancestors of the lobe-finned and ray-finned fishes.

In this study, we used micro-CT scans to examine the pectoral fins of *hoxaa^-/-^;hoxab^+/-^*;*hoxda^+/-^* and *hoxaa^+/-^;hoxab^+/-^*;*hoxda^-/-^* adult fish (Fig. 4). When comparing our findings to previously reported *hoxa13a;a13b* mutants and *hoxa13a;a13b;d13a* mosaic mutants (*hox13* mutants)^19^, we noticed both similarities and differences. One common feature is the increased number of distal radials, which is consistent across all these mutants. Additionally, both *hoxaa^-/-^;hoxab^+/-^*;*hoxda^+/-^* and *hoxaa^+/-^;hoxab^+/-^*;*hoxda^-/-^* fish exhibit morphological abnormalities in the posterior proximal radials, similar to *hox13* mutants^19^. Our multiple *hox* cluster deletion mutants retain some *hox13* genes, suggesting that other posterior *hox* genes in *HoxA-* and *HoxD-*related clusters may also play a role in the formation of proximal and distal radials. On the other hand, unlike *hox13* mutants where the fin rays are lost^19^, our mutants only show a slight reduction in the number of fin rays while still retaining the fin rays. This suggests that, unlike proximal and distal radials, a small amount of *hox13* may be sufficient for the formation of fin rays in the pectoral fin.

We also detected common abnormalities in proximal radials and fin rays in the posterior portion of the pectoral fins in *hoxaa^-/-^;hoxab^+/-^*;*hoxda^+/-^*and *hoxaa^+/-^;hoxab^+/-^*;*hoxda^-/-^* fish, (Fig. 4). As no apparent abnormalities were observed in these anterior skeletons, the reduced function of zebrafish *HoxA-* and *HoxD*-related clusters seems to preferentially influence the posterior portion of the pectoral fins. In zebrafish, the expression of posterior *HoxA-* and *HoxD*-related genes is progressively restricted to the posterior portion of the developing pectoral fins, after the formation of fin buds^18–20^. Furthermore, it has been shown that the phenotypes in which the posterior skeletons of the pectoral fin are altered by specific mutations can be suppressed by adding further null mutations in *hoxa11a*, *hoxa11b*, and *hoxd11a*^33^. Importantly, several lines of evidence have indicated that the posterior structure of the pectoral fin has a gene network similar to that of the limbs of tetrapods ^18,26–31^. Our study demonstrates that zebrafish *hox* clusters, homologous to *HoxA* and *HoxD* clusters, which are essential for limb development in mice, also play an important role in the specific skeletal structures of adult zebrafish pectoral fin which are considered to be potentially homologous to the tetrapod limbs.

In this study, zebrafish three *HoxA-* and *HoxD-*related clusters were shown to function together in zebrafish pectoral fin formation, similar to what has been observed in mice^8–15^. While the focus of this study was on their role in pectoral fin development, it is also possible that zebrafish *hoxaa*, *hoxab*, and *hoxda* clusters cooperate in other developmental processes. Interestingly, we have shown that mutants with more than two deletions of these clusters are lethal, although the exact cause of the lethality remains unknown. Some of the finless mutants in zebrafish can survive ^32,34,35^, suggesting that pectoral fin shortening does not necessarily result in lethality and may be linked to other developmental events related to survival. In mice, it has been shown that cooperative functions of *HoxA* and *HoxD* clusters are essential for urogenital development ^10^. This should be clarified in the future to elucidate the roles of *hoxaa*, *hoxab*, and *hoxda* clusters during zebrafish development.

## Materials and methods

### Zebrafish husbandry

Zebrafish (Riken WT; RW), which were provided by the National BioResource Project (NBRP) Zebrafish in Japan, were maintained at 27°C with a 14-h light/10-h dark cycle. Embryos were obtained from natural spawning and the larvae were raised at 28.5°C. Developmental stages of the embryos were determined based on days post-fertilization (dpf) and morphological features as previously described^36^. Zebrafish mutants used in this study are *hoxaa* cluster-deficient*^sud^*^111^, *hoxab* cluster-deficient*^sud^*^112^, *hoxda* cluster-deficient*^sud^*^116^, which were previously generated^21^.

### Genotyping of mutants

For the genotyping of mutants, PCR was carried out. To partially dissect the caudal fins in juvenile and adult fish, the fish were anesthetized using the minimum required dose while checking the response with tricaine. Then, genomic DNA was extracted by NaOH method and used as a PCR template ^37^. PCR was performed as described previously for the genotyping of *hox* cluster-deficient mutants^21^. After the reactions, the PCR products were separated by the electrophoresis in 2 % agarose gel in 0.5 x TBE buffer.

### Whole-mount *in situ* hybridization

Whole-mount *in situ* hybridization was performed for zebrafish embryos as previously described ^38^. After staining, the tail region was dissected with a disposable scalpel (Feather). Genotyping was performed using genomic DNA extracted from the dissected tail as described above. Stained embryos with the desired genotype were selected and photographed under a stereomicroscope (Leica M205 FA) with a digital camera (Leica DFC350FX).

### Alcian blue staining and the measurements of the endoskeletal disc and fin-fold

Zebrafish larvae at 5 dpf were anesthetized in tricaine or MS-222 and then fixed with 4 % paraformaldehyde in PBS. Alcian blue staining under strong acid conditions makes it difficult to extract the genomic DNA required for genotyping after staining. Thus, acid-free Alcian blue staining was performed as described previously^39^. After Alcian blue staining, genomic DNA was extracted from the posterior body, which was manually detached under a stereomicroscope with forceps, and the genotype was determined by PCR as previously described^21^. For the larvae with the desired genotype, pectoral fins were manually dissected with forceps under a stereomicroscope, mounted in 50 % glycerol on a glass slide (Matsunami), and photographed under a stereomicroscope (Leica M205FA). From these images, the longest lengths of the endoskeletal disc and fin-fold along the proximal-distal axis were measured using ImageJ software.

### X-ray micro-computed tomography (CT) scanning analysis

The pectoral fin skeletons of zebrafish were analyzed by X-ray micro-CT scanning as essentially previously described^40^. Briefly, adult zebrafish were anesthetized in 0.01% tricaine (w/v) and sacrificed in an ice water bath. Then specimens were fixed with 4 % paraformaldehyde in PBS at 4 °C overnight and transferred to 70 % ethanol.

The regions surrounding the pectoral fins were manually dissected by forceps from the zebrafish body under the stereomicroscope. Using an X-ray micro-CT (ScanXmate-E090S105; Comscantechno), the dissected pectoral fins were scanned at a tube peak voltage of 85 kV and a tube current of 90 µA. The micro-CT data were constructed using coneCTexpress software (Comscantechno) and stored as a dataset with an isotropic resolution of 3.1-3.4 µm. Three-dimensional image analysis was performed using OsiriX MD software (Pixmeo). Finally, movies were edited using Adobe Premiere Pro (Adobe) and DaVinci Resolve (Blackmagic Design).

### Digital dissection of zebrafish pectoral fin skeletal structures

Based on the data of the skeletal structures of the pectoral fin obtained from micro-CT scans, each zebrafish skeletal element was manually dissected using the Imaris contour surface feature (in the Measurement Pro) and displayed in a 3D image using Imaris v10.0 (Bitplane).

### Ethics approval

All the experiments using live zebrafish were approved by the Committee for Animal Care and Use of Saitama University (R2-A-1-6, R3-A-1-6, R4-A-1-7, R5-A-1-7) and carried out according to the Animal Research Reporting of In Vivo Experiments (ARRIVE) guidelines and to relevant guidelines and regulations.

## Supporting information

Ishizaka et al-Supplementary Data

## Acknowledgments

We thank the NBRP zebrafish for providing the RW strain and for preserving zebrafish *hox* cluster-deficient lines used in this study. We also thank Koji Tamura and Gembu Abe for providing the zebrafish *tbx5a* plasmid for *in situ* hybridization. This work was supported by KAKENHI Grants-in-Aid for Scientific Research from the Ministry of Education, Culture, Sports, Science, and Technology, Japan (18K06177, 23K05790 to A.K.) and by the National Institute of Genetics under the Joint Research and Research Meeting (NIG-JOINT) program (38A2019, 7A2020, 66A2021, 18A2022, 31A2023 to A.K.).

## Data availability

The datasets used and/or analysed during the current study are available from the corresponding author on reasonable request.

## Author Contributions

A.K. conceived and designed the experiments, M.I., H.N., H.K., R.F., T.T., S.O., R.K., and A.K. performed the embryonic phenotype analysis, A.M. performed ray micro-CT scan analysis and constructed the 3D-movie, A.K. wrote the manuscript. All authors discussed the results and commented on the manuscript.

