## Supplementary material for "The functional roles of zebrafish *HoxA*- and *HoxD*-related clusters in the pectoral fin development": Ishizaka et al-Supplementary Data

**Supplementary data by Ishizaka et al.**

**Figure S1**. **Schematic representation of *Hox* genes in zebrafish and mouse *Hox* clusters.**

Zebrafish possess 49 *hox* genes spread across seven *hox* clusters. Mice possess 39 *Hox* genes in four *Hox* clusters.

**Figure S2. Quantification of *shha* signals in the pectoral fin at 48 hpf embryos.**

Using images of *in situ* stained embryos, the *shha* signals in the developing pectoral fin were measured for both the left and right pectoral fins of three individuals for each genotype. Measurements was carried out using ImageJ, where the area of *shha* expression in the pectoral fin and its intensity (mean) in the region were recorded. The same sized area of unstained yolk was also measured as background, and the difference between the two values was calculated. The area and intensity values were then multiplied to obtain the expression level of *shha*, and the relative expression level was presented as a graph with the mean value of the wild-type set to 100%. Statistical analysis was conducted using the Tukey-Kramer test with $^{**}P < 0.01$ and $^{****}P < 0.0001$. The error bars represent the standard error of the mean.

**Movie S1. Magnified skeletal structures of the pectoral fin in adult wild-type zebrafish.**

Micro-CT scan was performed to magnify the skeletal structures of the pectoral fin in adult wild-type zebrafish. The left pectoral fin is displayed. A 3D movie was created using OsiriX MD ver13. Scale bar: 500 μm.

**Movie S2**. **Magnified skeletal structures of the pectoral fin in adult *hoxaa$^{-/-}$;ab$^{+/-}$;da$^{+/-}$* zebrafish.**

Micro-CT scan was performed to magnify the skeletal structures of the pectoral fin in adult *hoxaa$^{-/-}$;ab$^{+/-}$;da$^{+/-}$* fish. The left pectoral fin is displayed. A 3D movie was created using OsiriX MD ver13. Scale bar: 500 μm.

**Movie S3**. **Magnified skeletal structures of the pectoral fin in adult *hoxaa*$^{+/-}$*;ab*$^{+/-}$*;da*$^{-/-}$ zebrafish.**

Micro-CT scan was performed to magnify the skeletal structures of the pectoral fin in adult *hoxaa*$^{+/-}$*;ab*$^{+/-}$*;da*$^{-/-}$ fish. The left pectoral fin is displayed. A 3D movie was created using OsiriX MD ver13. Scale bar: 500 μm.

**Movie S4**. **CT segmentation of skeletal structures of the pectoral fin in adult wild-type zebrafish.**

3D movie showing proximal radials (gray) and distal radials (red) of the pectoral fin in in wild-type fish. The left pectoral fin of the same individual shown in Movie S1 was used. A 3D movie was created using Imaris ver10.

**Movie S5**. **CT segmentation of skeletal structures of the pectoral fin in adult *hoxaa*$^{-/-}$*;ab*$^{+/-}$*;da*$^{+/-}$ fish.**

3D movie showing proximal radials (gray) and distal radials (red) of the pectoral fin in *hoxaa*$^{-/-}$*;ab*$^{+/-}$*;da*$^{+/-}$ fish. The left pectoral fin of the same individual shown in Movie S2 was used. A 3D movie was created using Imaris ver10.

**Movie S6**. **CT segmentation of skeletal structures of the pectoral fin in adult *hoxaa*$^{+/-}$*;ab*$^{+/-}$*;da*$^{-/-}$ fish.**

3D movie showing proximal radials (gray) and distal radials (red) of the pectoral fin in *hoxaa*$^{+/-}$*;ab*$^{+/-}$*;da*$^{-/-}$ fish. The left pectoral fin of the same individual shown in Movie S3 was used. A 3D movie was created using Imaris ver10.

**Movie S7**. **2D movie showing the proximal radials and distal radials in wild-type, *hoxaa*$^{-/-}$*;ab*$^{+/-}$*;da*$^{+/-}$, and *hoxaa*$^{+/-}$*;ab*$^{+/-}$*;da*$^{-/-}$ fish.**

The 2D movie display the proximal radials (yellow) distal radials (magenta), and fin rays (cyan) in wild-type, *hoxaa*$^{-/-}$*;ab*$^{+/-}$*;da*$^{+/-}$, and *hoxaa*$^{+/-}$*;ab*$^{+/-}$*;da*$^{-/-}$ fish. These outlines were used to create Movies S4-S6. A 3D movie was created using Imaris ver10.

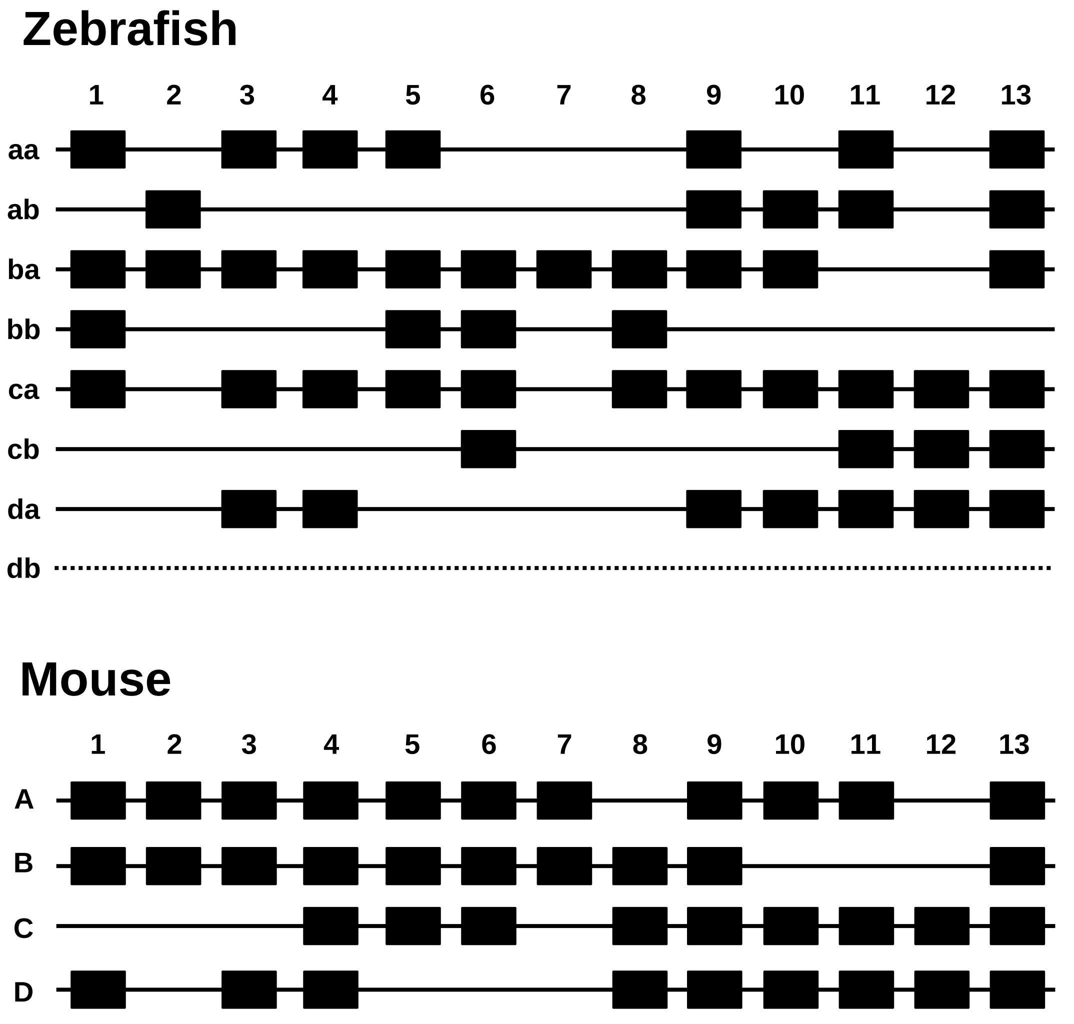


**Fig. S1**. **Schematic representation of *Hox* genes in *Hox* clusters in zebrafish and mouse.**

Zebrafish possess 49 *hox* genes distributed across seven *hox* clusters. Mice possess 39 *Hox* genes in four *Hox* clusters.

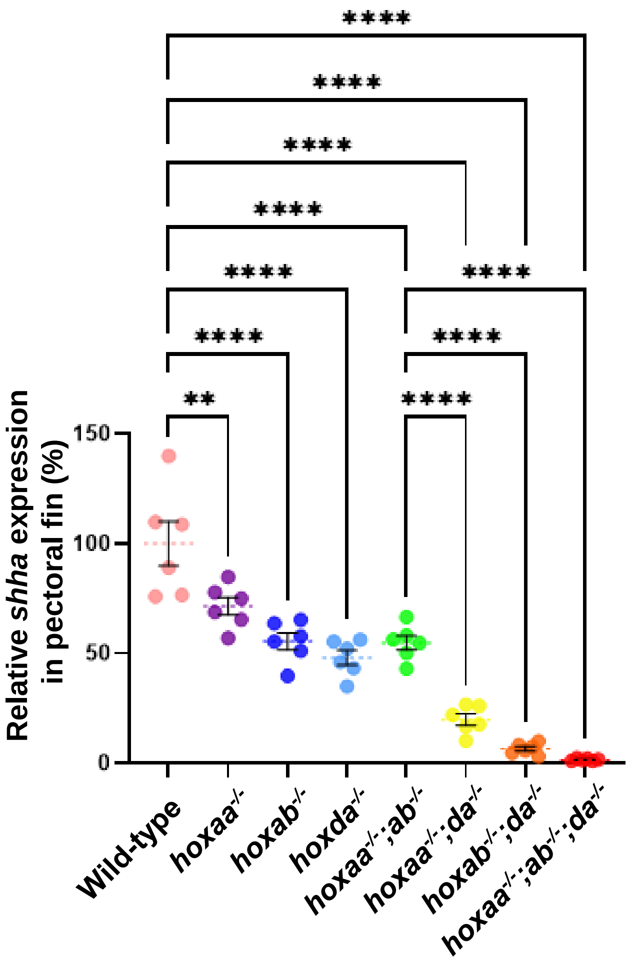


**Fig. S2. Quantification of *shha* signals in the pectoral fin at 48 hpf embryos.** Using images of the *in situ* stained embryos, the *shha* signals in the developing pectoral fin of both the left and right pectoral fins of the three individuals for each genotype were measured by ImageJ. The area of *shha* expression in pectoral fin and the intensity (mean) in the region were measured. The same size of the area of unstained yolk was also measured as background and the difference between the two was calculated. The area and intensity (mean) values were multiplied to obtain the expression level of *shha* expressions, and the relative expression level was shown as a graph with the mean value of wild-type as 100%. The Tukey-Kramer test was performed with $**P < 0.01$ and $****P < 0.0001$. Error bars represent the standard error of the mean.
